# deMULTIplex2: robust sample demultiplexing for scRNA-seq

**DOI:** 10.1101/2023.04.11.536275

**Authors:** Qin Zhu, Daniel N. Conrad, Zev J. Gartner

**Affiliations:** University of California San Francisco, Department of Pharmaceutical Chemistry, San Francisco, CA 94158; Chan Zuckerberg Biohub, University of California San Francisco, San Francisco, CA 94158; Center for Cellular Construction, University of California, San Francisco, CA 94158

## Abstract

Single-cell sample multiplexing technologies function by associating sample-specific barcode tags with cell-specific barcode tags, thereby increasing sample throughput, reducing batch effects, and decreasing reagent costs. Computational methods must then correctly associate cell-tags with sample-tags, but their performance deteriorates rapidly when working with datasets that are large, have imbalanced cell numbers across samples, or are noisy due to cross-contamination among sample tags - unavoidable features of many real-world experiments. Here we introduce deMULTIplex2, a mechanism-guided classification algorithm for multiplexed scRNA-seq data that successfully recovers many more cells across a spectrum of challenging datasets compared to existing methods. deMULTIplex2 is built on a statistical model of tag read counts derived from the physical mechanism of tag cross-contamination. Using generalized linear models and expectation-maximization, deMULTIplex2 probabilistically infers the sample identity of each cell and classifies singlets with high accuracy. Using Randomized Quantile Residuals, we show the model fits both simulated and real datasets. Benchmarking analysis suggests that deMULTIplex2 outperforms existing algorithms, especially when handling large and noisy single-cell datasets or those with unbalanced sample compositions.

## Background

Single cell sequencing has revolutionized biomedical research by providing an unbiased, high-resolution, and high-throughput profile of healthy and diseased tissues (Svensson et al., 2018). Recent advances in single-cell sample multiplexing technologies, such as those based on lipid-tagged indices (McGinnis, Patterson, et al., 2019), barcoded antibodies (Gaublomme et al., 2019; Mimitou et al., 2019; Stoeckius et al., 2018), chemical labeling (Gehring et al., 2020), Nuclear hashing (Srivatsan et al., 2020), lentiviral infection (Guo et al., 2019), transient transfection (Shin et al., 2019), and genetic variation (Heaton et al., 2020; Huang et al., 2019; Kang et al., 2018; McFarland et al., 2020), further improves the scalability of scRNA-seq, allowing multiple samples from different experimental condition to be pooled together and sequenced. These procedures greatly reduce experimental costs and batch effects while increasing cell throughput, but require demultiplexing of the tag count data to assign each cell to the correct sample-of-origin. In an ideal experiment, cells from a sample will be uniquely labeled by only a single tag, and subsequent demultiplexing based on the tag count would be trivial. In reality, however, ambient or debris-bound lipid- and cholesterol-modified oligos (LMO/CMOs) or antibody-derived tags (ADTs) (both referred to here as “tags”) may bind to or co-encapsulate with cells from other samples when pooled. These contaminating tags, along with variation in tag capture rate and the inherent technical noise of single cell sequencing technology, manifest in real data as large numbers of off-target tags (noise) associated with each cell in addition to the on-target tags (signal). The signal-to-noise ratio can vary significantly between cell types and samples, complicating the essential task of identifying a clean cut-off between cells from different samples.

To address this challenge, several computational approaches have been implemented. The deMULTIplex R package, which was released together with MULTI-seq (McGinnis, Patterson, et al., 2019), assumes that the positive and negative cells for each tag follow a bimodal distribution, and uses local maxima of a smoothed probability density function (PDF) and quantile sweep to define the threshold for each tag. Similar to the bimodal distribution assumption, GMM-Demux (Xin et al., 2020) fits a Gaussian mixture model to the tag count data, and uses Bayesian estimation to determine the sample identity of each cell. BFF is another method which was developed based on the bimodal distribution assumption, and offers two modes of classification, one based on raw count (BFF_raw_) and one based on normalized counts (BFF_cluster_). The HashedDrops function in the R package DropletUtils (Lun et al., 2019) is a straightforward method which assigns each cell to a sample based on its most abundant tag, and uses the log fold change between the highest and second-highest tag counts to represent the confidence of assignment. The HTODemux function in the Seurat package first clusters cells in the tag count space, and then uses the cluster with lowest average tag abundance to fit a negative binomial distribution to define the threshold of calling positive cells (Stoeckius et al., 2018). DemuxEM first estimates the background count distribution using empty droplets, then applies the expectation-maximization (EM) algorithm to determine the fraction of a cell’s tag signal coming from the background or the true staining, and performs classification on the background-subtracted data (Gaublomme et al., 2019). Lastly, a recent method, demuxmix, uses regression mixture models to account for the positive association between tag count and the number of detected genes, leading to improved classification (Klein, 2023). These algorithms all rely on a specific statistical feature of the tag count distribution, such as the bimodal distribution, the enrichment of tag in positively labeled cells, or the association between tag count and gene count, to identify a decision boundary in the relevant feature space. However, these assumptions do not account for the fundamental physical mechanisms through which distinct tag distributions arise across droplet-based scRNA-seq data. As a consequence, they fail when basic assumptions are not met—such as when sample composition is unbalanced, or tag cross-contamination is high.

Here, we introduce deMULTIplex2, which models tag cross-contamination in a multiplexed single cell experiment based on the physical mechanism through which tag distributions arise in populations of droplet-encapsulated cells. We first derive the analytical form of the expected tag count, and show that for each tag, the count distribution can be modeled with two negative binomial generalized linear models (GLM-NB) in two distinct spaces. Using EM, we robustly fit the two GLM-NBs in the corresponding space and probabilistically determine if a cell is positively labeled by a tag. The distribution of randomized quantile residuals (RQR) suggests that this model fits well on both simulated data and real data with different degrees of noise. When benchmarking deMULTIplex2 against existing methods, we were able to classify significantly more cells with high precision, and the method performs consistently well on noisy, large-scale scRNA-seq datasets generated with diverse multiplexing technologies.

## Results

### Modeling tag cross-contamination

During a single cell multiplexing experiment, cells are first incubated (labeled) with a sample-specific tag, and then pooled together for single-cell capture and sequencing. The initial labeling of cells in each sample may have variable conditions (i.e., tag concentration, staining time, debris which sequesters tags, etc.), but the contamination happens only after pooling when all cells are in the same solution. Therefore, we focus on modeling the contamination process post pooling, which we assume occurs under the same conditions across all the cells regardless of which sample they came from because all cells are bathed in the same buffer solution.

Consider a simple experiment consisting of two samples, labeled with tags A and B respectively (Fig. 1A). After the samples are pooled together, excess tag B can bind to cells that were initially labeled with tag A (tagA+/tagB− cells) and vice versa, causing contamination. We model this process as a simple chemical reaction between the cell surface S and tag B:

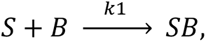

where *k*1 is the rate constant of the reaction. Typically, the reaction will not reach equilibrium because there will be limited incubation time at low temperature prior to single cell capture (as recommended by most protocols). Therefore, the “concentration” of cell-bound tag B, as denoted by [SB], can be expressed as

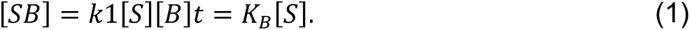

**Figure 1.**
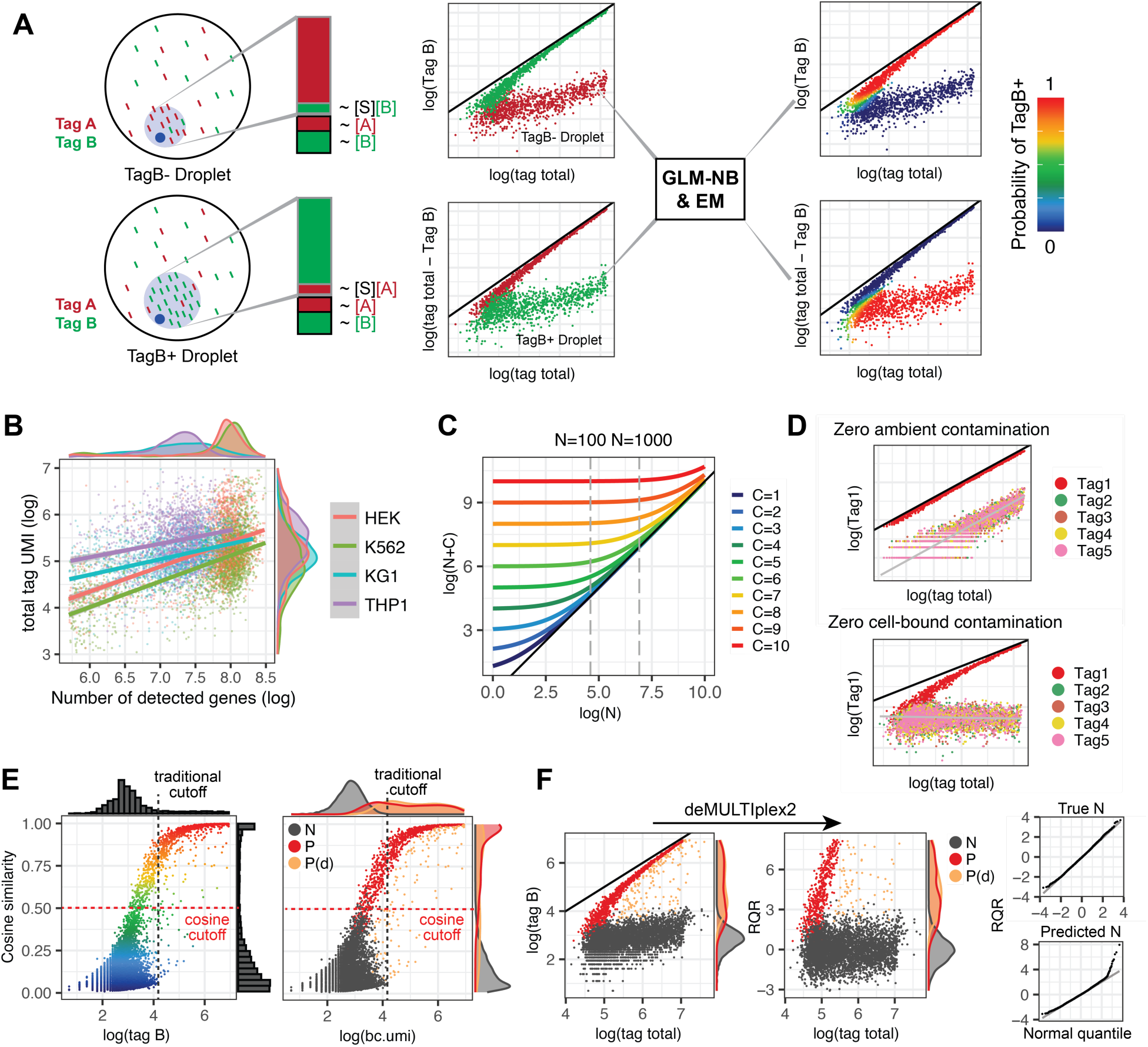
An overview of the deMULTIplex2 algorithm. (A) Illustration for how deMULTIplex2 models tag cross-contamination in a simple two-sample multiplexing experiment. Each sample is labeled with tag A or B. Pooling the samples for single cell capture allows floating tags to be bound to the cell or captured by the droplet volume. These contaminating tag counts are modeled by fitting two GLM-NB models in two separate spaces, and the sample identity is inferred using the EM algorithm. The y=x line is shown in black for this panel and panels C,D,F. (B) Scatter plot illustrating the association between total tag UMI count and number of detected genes in the 4-cell-line dataset from Stoeckius et al (Stoeckius et al., 2018). (C) Plot of the relationship between log(N+C) and log(N) across different values of C. For a typical experiment, the total tag count of cells is enriched within a range of one to two orders of magnitude (such as in the range of 100 to 1000 highlighted by the dashed line). (D) Simulated tag count distribution for scenarios with zero ambient contamination or zero cell-bound contamination. (E) Cosine similarity with canonical vector plotted against the tag count for a given tag (tag B) using simulated data. Several existing methods look for a bimodal distribution in the count dimension, while deMULTIplex2 takes advantage of the separation between positive and negative cells in the cosine dimension to initialize EM. (F, left) The original UMI count of a given tag plotted against total tag count with simulated data. (F, right) RQRs computed by deMULTIplex2 of the same simulated data plotted against total tag counts. N: negative cells, P: positive cells, P(d): doublets positive with tag B. The RQRs are plotted against the normal quantiles for true negative cells and predicted negative cells.

*K_B_* is assumed to be a constant which is uniform across all the cells because of the same binding mechanism (*k*1), ambient concentration ([*B*]), and incubation time (*t*).

This suggests that the final bound tag count is proportional to the total cell surface area, and larger cells tend to get more bound tags. Indeed, when plotting the total number of bound tags against the total number of genes (which is typically considered to be correlated with cell size), positive correlations are observed across different cell types (Fig. 1B) (Stoeckius et al., 2018). The demuxmix method builds upon this observation to fit a regression mixture model with total detected gene count as a predictor to account for the extra variance observed in tag count data (Klein, 2023). However, gene count is highly cell-type-specific and can lead to cell-type-biased classifications (Fig. 1B). On the other hand, the total tag counts exhibit much less cell-type specificity, and therefore may be more associated with cell surface area (Fig. 1B), i.e., *N_total_* ∝ [*S*]. This additional assumption allows us to re-write equation (1) to represent the expected tag count of contaminating B as a fraction of total tag counts:

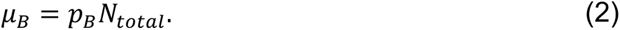

Due to sampling variation inherent to scRNA-seq technology, the observed unique molecular identifier (UMI) count is commonly modeled with a Poisson or Negative binomial (NB) distribution (Hafemeister & Satija, 2019). In practice, we and others have observed over-dispersion in the tag UMI count data (Klein, 2023; Stoeckius et al., 2018). We therefore choose to use a negative binomial distribution and fit the observed tag counts *X_B_* with a generalized linear model:

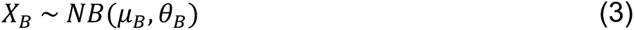

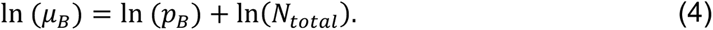

Equations (2-4) are similar to a proposed analytical solution of scRNA-seq UMI count distributions (Lause et al., 2021). As Lause et al. pointed out, this model suggests that the linear coefficient *β*_1_ before ln (*N_total_*) should be fixed to 1 for negative control cells without biological variability (in this case, cells that have not been labeled with the contaminating tag prior to pooling). However, although the fit of *β*_1_ for some real datasets is indeed very close to 1, the majority of datasets we have tested demonstrate an estimated *β*_1_ lower than 1 (Fig. S1). When inspecting the count distributions of these datasets, however, we realized that there is a second source of contamination, where ambient floating tags or debris-bound tags are co-encapsulated in the droplet and get sequenced along with the cell-surface-bound tags (Fig. 1A, Fig. S1). Under such a model, the expected UMI count of ambient B (denote as *M_B_*) is constant across cells assuming consistent droplet size, and is proportional to the concentration of ambient B. Combining these two sources of contaminating tags, equation 2 for the expected count of contaminating tag B should be revised as:

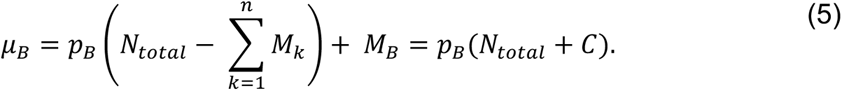

Note that 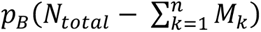 is representing the cell-bound contamination previously defined by Equation 2, but with the total cell-bound tag count re-calculated by excluding the sum of all ambient tag count (*M_k_* over *n* tags) from the total observed tag count. 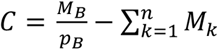, which is the same across all negative cells contaminated with tag B and does not depend on the total observed tag count, thus can be treated as a constant. Therefore,

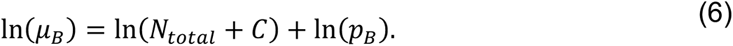

In the above equation, the relationship between ln(*μ_B_*) and ln(*N_total_*) is no longer linear. *C* is a tag-specific constant which is difficult to estimate. However, looking at the relationship between ln(*N_total_* + *C*) and ln(*N_total_*) across different values of *C*, we found within a limited range of ln(*N_total_*), such as that typically observed in total tag counts (one or two orders of magnitude of difference in total tag count, likely limited by the size range of eukaryotic cells, Fig. 1B), their relationship is approximately linear (Fig. 1C), i.e.,

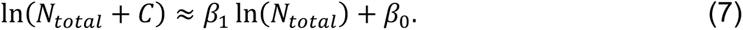

Then

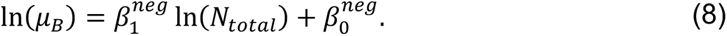

Here, 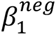 and 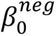 are linear coefficients that can be estimated with a GLM-NB model. Modeling these two sources of contamination allows us to simulate datasets with different ratios of cell-bound and ambient tag contamination. Encouragingly, simulated data qualitatively reproduces a variety of distributions we see in real datasets (Fig 1D, Fig S1).

To specify the full probabilistic model for the data, we also need to model the count distributions of the positive cells which were originally labeled with tag B. As discussed before, we cannot directly model the count of tag B prior to pooling. But in the simple experiment illustrated in Fig. 1A, positive cells of B are also the negative cells of A, meaning that the same GLM-NB model described by Equation 3 and 8 can be applied to the tag count of A (*X_A_* = *N_total_* − *X_B_*) to model the distribution of the positive cells of B.

More generally, for the positive cells labeled with a particular tag B, we consider the distribution of total contamination count *N_total_* − *X_B_*, where

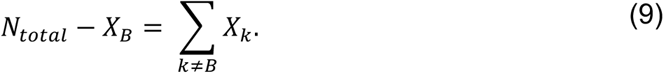

Assuming the counts of each of the contaminating tags follow a NB distribution, the total contamination is the convolution of multiple NB distributions, which also has the form of NB distribution having a mean equal to the sum of all means of contaminating tags:

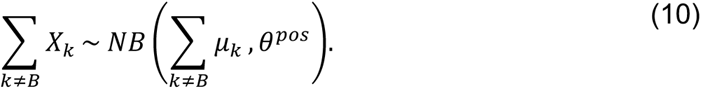

This result makes intuitive sense because the tags share the same chemical and physical properties, so the pool of contaminating tags can be thought as a single contaminating meta-tag. Following derivation similar to that for single-tag contamination, the expected tag count of multi-tag contamination follows:

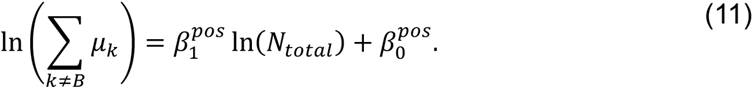

We provide the detailed derivation of (11) in **Methods**. Equations 9–11 suggests that the distribution of positive cells can be modeled with a second GLM-NB model in the *N_total_* − *X* vs *N_total_* (total tag count minus observed count of positive tag vs. total tag count) space (Fig. 1A). It is important to point out that the distribution of positive cells in the *X* vs *N_total_* space (positive tag count vs total tag count) is non-linear with ambient contamination. The cells converge to the y=x line with increased signal-to-noise ratio, but can never cross the y=x line (Fig. 1A, D). Therefore, regression mixture models, such as that proposed by demuxmix, cannot properly fit positive cells in this space, and will likely result in poor classification when ambient contamination is present.

### Probabilistic classification of cells with expectation-maximization

The two GLM-NB models specified in the separate spaces allow us to define the joint probability distribution of all cells and use expectation-maximization (EM) to solve for the identity (positive or negative) of each cell for each tag. The joint probability distribution for each tag can be expressed as:

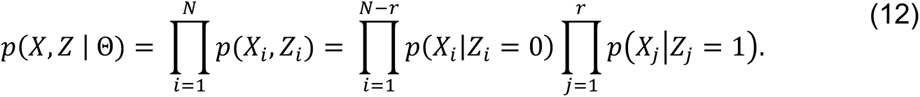

Here, *N* is the total number of cells, *r* is the number of positive cells, and the latent variable *Z_i_* indicates whether a cell *i* is positively labeled by the tag. For each tag (T) and each cell, the conditional probability follows the NB distribution derived in the previous section, i.e.,

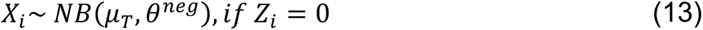

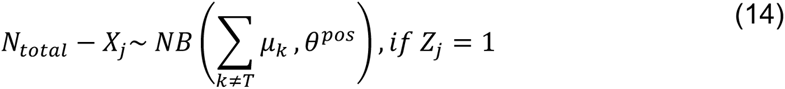

The EM algorithm iterates between estimating each cell’s identity, as described by *Z*, and fitting the two GLM-NB model in the corresponding spaces. The algorithm stops upon convergence, or when the user-specified maximum number of iterations has been reached. In practice, we found that for most datasets the algorithm quickly converges within a few iterations when using a reasonable number of cells for model fitting (Fig. S2). Because random sampling of a few thousand cells is enough for robust fitting of the GLM-NB models (Fig. S2), deMULTIplex2 can process large datasets with high speed, low memory requirement, and robust performance.

Finally, upon convergence, deMULTIplex2 reports the posterior probability of a cell being positively labeled by each tag (**Methods**, Fig. 1A). The decision boundary produced by the algorithm usually has a large margin, with relatively small difference in assignment results from different choice of probability cutoff. Therefore, deMULTIplex2 can resolve each cell’s identity with high confidence in a probabilistic manner. With each cell being classified as positive or negative for each tag, deMULTIplex2 determines whether a cell is a singlet, a multiplet (generally referred to as “doublet”) or negative (not labeled with any tag) based on the total count of positive tags.

### Initializing EM with cosine similarity cutoff

The EM algorithm is known to be sensitive to initialization (Meila & Heckerman, 2013; Melnykov & Melnykov, 2012). Previous efforts have addressed this issue through multiple randomly initialized short runs (Baudry & Celeux, 2015; Biernacki et al., 2003; Grun & Leisch, 2008) or through an initial clustering (Fraley & Raftery, 2006; Klein, 2023). However, these strategies require additional computation and may still fail due to imbalanced cell number between positive cells and negative cells (which is typical for a multiplexed dataset). Therefore, we sought a statistic, derived from the unique features of positive cells and negative cells, that robustly generates a satisfactory initial separation among positive and negative cells and can properly initialize the EM algorithm.

We found that the cosine similarity between the tag count vector of each cell and the canonical vectors for each tag (i.e., a vector withs 1 and 0s, where position of 1 indicate which tag the vector represents) provides a close-to-truth initial guess for the identity of each cell. The cosine metric can be understood using a barnyard plot. Assuming low contamination in a two-tag mixture experiment, true positive singlets will be aligned with each axis; the resulting cosine similarity with the canonical vectors <1,0> and <0,1> will be 1 and 0 or vice versa. In real datasets where true positive tags take up the majority of the tag reads in a cell, the cosine similarity, when plotted against the tag count, approximately follows a sigmoid curve (Fig. 1E). The distribution in the tag count dimension is traditionally used to define a cutoff for positive vs negative cells and its bimodality is the core assumption of many existing methods. However, in experiments with many pooled samples, imbalanced numbers of cells per sample, or high background noise, the second positive peak could be undetectable or could overlap significantly with the negative peak (Fig. 1E), leading to failures in methods that rely heavily on the assumption of bimodality. The cosine similarity, in contrast, provides a second dimension with larger margins for drawing the initial cutoff to initialize the EM (Fig. 1E). It correctly enriches for true positive cells on one side of the sigmoid, and for true negative cells on the other side. With both real data and simulated data, we found that EM initialized with a cosine cutoff in the range of 0.2 to 0.9 quickly and robustly converges to the correct fit (Fig. S2).

### Randomized quantile residuals for diagnosing the goodness-of-fit

Residual plots show the discrepancy between data and model and are commonly used to diagnose goodness-of-fit. In a normal linear model, the Pearson residual, defined as 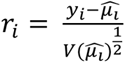, is asymptotically normal under the true model. However, for generalized linear models for Poisson or NB distributions, the residual is far from normal due to the discrete response values. Randomized quantile residuals (RQRs) were proposed (Dunn & Smyth, 1996) to overcome this problem and have been applied for diagnosing GLM models for count data (Bai et al., 2021; Feng et al., 2020). RQR is an extension of the quantile residual (QR), which inverts the fitted distribution function of each observation to the corresponding normal quantile. QR is defined as:

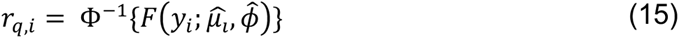

To generalize QR to the discrete cumulative distribution F of Poisson and NB, a random uniform sampling was performed for each observation to obtain a continuous mapping to the normal distribution, i.e.,

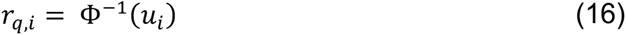

where *u_i_* is drawn randomly from a uniform distribution defined on the interval 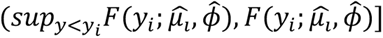.

Therefore, under the null hypothesis, the RQR of a well-specified model will be normally distributed.

Using RQR, we evaluated the goodness-of-fit of the two GLM-NB models for the positive cells and negative cells using both simulated data and real data. We plotted the RQRs against the standard normal quantiles, also known as the Q-Q plot, for each regression fit. We found that for true negative cells, RQRs are indeed normally distributed, and for deMULTIplex2-predicted negative cells, the RQRs are very close to normal, and are often right-skewed due to the ambiguity at the boundary of positive and negative cells (Fig. 1F, S1). For positive cells in the *N_total_* − *X* vs *N_total_* space, however, the RQRs deviate from normal for several tags in the real dataset (Fig. S1), likely due to the presence of doublets. Interestingly, we found the final classification are still very close to the ground truth despite this imperfect fit, likely because the distinct distribution of positive and negative cells in the two spaces led to large difference between predicted positive probability and negative probability from the two GLM models, and any misclassification during the EM will incur a high cost and will be corrected in the next few EM iterations.

### deMULTIplex2 outperforms other methods on simulated data

The two-component contamination model allows us to simulate experimental datasets which recapitulate tag distributions we see in real datasets. We can therefore use these simulated data to evaluate the robustness and sensitivity of demultiplexing algorithms to key factors that affect classification accuracy, such as degree of contamination and sample size.

We first simulated a small 5-tag dataset with a mild degree of ambient and cell-bound contamination and a 10% doublet rate. The resulting UMAP based on the raw tag UMI count resembles what we often see from real datasets, with ambiguous, low-tag-count cells in the center and high-tag-count cells at the periphery. deMULTIplex2 was able to correctly predict the label of 4381 out of 5000 simulated singlets (excluding homotypic doublet) (Fig. 2A,B). When plotting the UMAP based on deMULTIplex2-computed RQRs, the embedding is much less influenced by the total tag count, and the doublets are placed at the periphery of each cluster, suggesting that RQR provides a proper normalization of the tag count data in addition to its diagnostic capability. The average precision of deMULTIplex2 is comparable to other methods, and the recall of deMULTIplex2 is second to the best-performing method, BFF_cluster_ (Fig. 2C).

**Figure 2.**
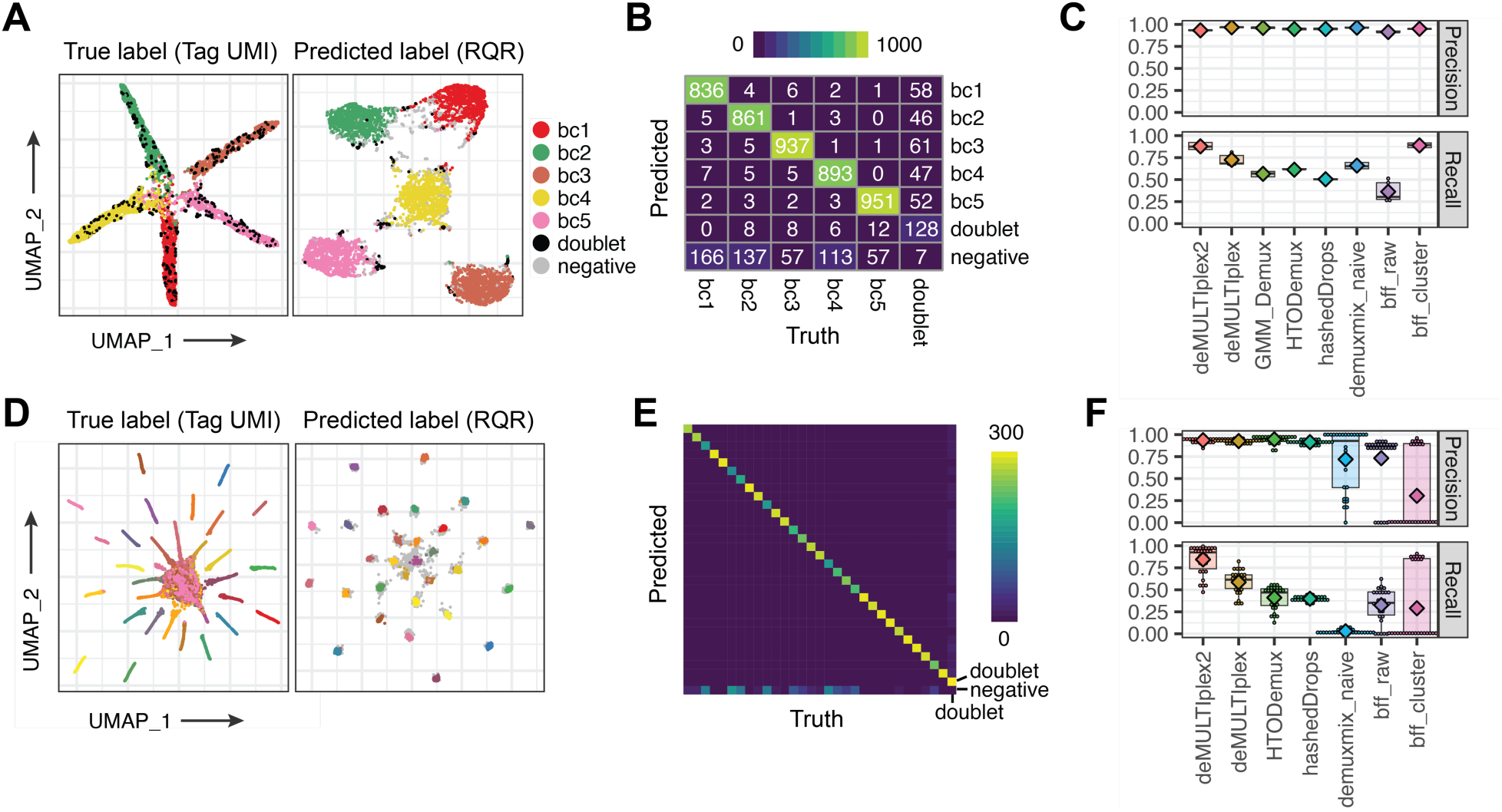
Performance of deMULTIplex2 on simulated datasets. (A-C) deMULTIplex2 results from a 5-tag simulated dataset with 1000 singlets per sample and 10% doublet rate. (D-F) deMULTIplex2 results from a 30-tag simulated dataset with 300 singlets per sample and a 10% doublet rate. (A) UMAP computed with raw tag UMI count or deMULTIplex2-computed RQRs. (B) Heatmap displaying the concordance between deMULTIplex2 results and true sample identities. Note that homotypic doublets are treated as singlets and “doublet” refers to heterotypic doublets. (C) Per-tag performance of deMULTIplex2 compared to other methods. Mean values are highlighted with the diamond points. Methods that require mRNA count matrix as input were excluded from this comparison. (D) UMAP computed with raw tag UMI count or deMULTIplex2-computed RQRs for the 30-tag simulated dataset. Doublets and negative cells are all colored grey for clean visualization. (E) Heatmap displaying the concordance between deMULTIplex2 results and true sample identities. (F) Per-tag performance of deMULTIplex2 compared to other methods. Mean values are highlighted with the diamond points.

We next asked how these algorithms perform on larger-scale datasets. We simulated a 30-tag, 300-cell-per-tag dataset with varying degree of contamination for each tag and a 10% doublet rate. The ambient contamination level is much higher for some tags, causing ambiguous cells from these samples to mix in the center of the raw-tag-count-based UMAP. Encouragingly, deMULTIplex2 was able to salvage many real singlets from these high-contamination samples (Fig. 2D,E), and the RQR-based UMAP shows a well-resolved structure. In contrast, the performance of other methods quickly deteriorates with increased data scale, and the best-performing method on the small dataset – BFF_cluster_ – becomes one of the worst-performing methods (Fig. 2F). The naïve mode of demuxmix also failed to classify most of the singlets, suggesting a simple regression mixture model cannot fully capture the distribution of the tag counts. In comparison, deMULTIplex2 achieved the second highest average precision (0.936) and highest average recall (0.844). HTOdemux has the highest average precision (0.946), but much lower average recall (0.410), mainly because the method was only classifying the high-count, high-confidence cells. In theory, achieving high recall without sacrificing precision is much more difficult because higher recall requires the method to draw decision boundary closer to the ambiguous and negative territory (without crossing the border to incur misclassification). In practice, high recall is often desired because it means many more real singlets can be recovered from expensive single cell experiments. In this case, deMULTIplex2 was able to correctly classify 7600 out of 9000 real singlets, while HTOdemux was only able to correctly classify 3696 cells, about half of that of deMULTIplex2.

### deMULTIplex2 outperforms other methods on real-world datasets

We assembled nine real-world datasets with associated ground-truth information to benchmark the performance of deMULTIplex2 and other methods. These datasets include the 8-donor-PBMC MULTI-seq and SCMK dataset from McGinnis et al., 2021 (McGinnis et al., 2021), the 4-cell line and 8-donor PBMC datasets from Stoeckius et al., 2018 (Stoeckius et al., 2018), the 8-donor single nucleus human cortex datasets from Gaublomme et al., 2019 (Gaublomme et al., 2019), the three batches of multi-donor bronchoalveolar lavage (BAL) datasets from Howitt et al., 2022 and Maksimovic et al., 2022 (Howitt et al., 2022; Maksimovic et al., 2022) and the human lung cell line dataset from Howitt et al., 2022 (Howitt et al., 2022). On datasets collected from different donors, SNP-based classification was used to obtain the ground-truth labels (Heaton et al., 2020; Huang et al., 2019). For datasets comprising different cell lines, ground truth labels were obtained by clustering in the transcriptomic space. In addition to precision and recall, we also report F-score (harmonic mean of precision and recall) as a balanced statistic for overall performance. As shown in Fig. 3A and Table S1, deMULTIplex2 consistently demonstrated superior performance in singlet classification, while its performance on doublet calling is comparable to existing methods (Fig. S3). Among these datasets, the MULTI-seq dataset from McGinnis et al., 2021 (McGinnis et al., 2021), and the ADT datasets from Stoeckius et al., 2018 (Stoeckius et al., 2018) and Gaublomme et al., 2019 (Gaublomme et al., 2019) are clean datasets with a very low degree of contamination. Therefore, all methods were able to achieve high precision with only a few exceptions. Notably, deMULTIplex2 was able to achieve highest average recall for most of these datasets, suggesting it can robustly retrieve real singlets without sacrificing precision (Table S1).

**Figure 3.**
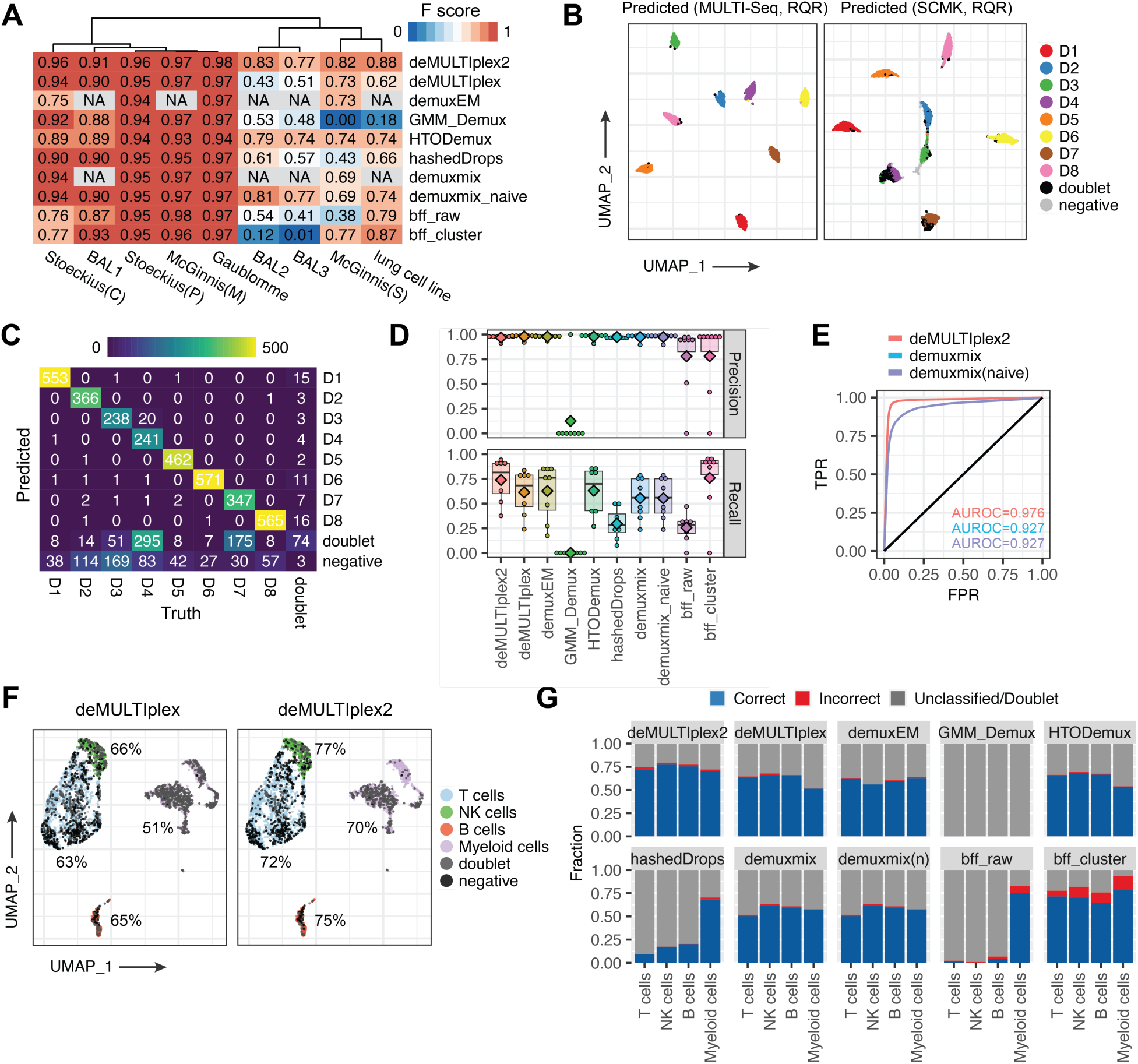
Performance of deMULTIplex2 on real datasets. (A) Heatmap summarizing the F score of deMULTIplex2 and other methods on 9 real datasets. Stoeckius(C) and Stoeckius(P) are cell line and multi-donor PBMC datasets from Stoeckius et al. (Stoeckius et al., 2018). McGinnis(M) and McGinnis(S) are MULTI-seq and SCMK datasets from McGinnis et al (McGinnis et al., 2021). BAL1, 2, 3 are three batches of multi-donor bronchoalveolar lavage (BAL) datasets from Howitt et al., 2022 and Maksimovic et al., 2022 (Howitt et al., 2022; Maksimovic et al., 2022). The lung cell line dataset is also from Howitt et al., 2022 (Howitt et al., 2022). NA indicates the method cannot be run on the corresponding datasets due to the unavailability of mRNA count matrix or an error (i.e., demuxEM returns an error on the SCMK dataset). (B) UMAP computed with deMULTIplex2-computed RQR for the MULTI-seq and SCMK datasets from McGinnis et al. (McGinnis et al., 2021), colored by donor ID predicted by deMULTIplex2. (C) Concordance between deMULTIplex2-predicted donor ID and the true donor ID based on SNP-based sample classification using souporcell (Heaton et al., 2020). (D) Performance of deMULTIplex2 and other methods on each sample. Mean values are highlighted with the diamond points. (E) Multiclass ROC curve of deMULTIplex2 and the two modes of demuxmix. False positive rate (FPR) and true positive rate (TPR) were computed for all samples using a one-vs-rest scheme and averaged to generate the ROC curve. (F) deMULTIplex and deMULTIplex2 recovered cells in the gene expression space. Percentage of correctly classified singlets are highlighted for each of the cell type. (G) Classification accuracy of each cell type across methods.

The BAL dataset consists of three separate batches with batch 2 and 3 having higher levels of contamination and doublet rates compared to batch 1 (Howitt et al., 2022; Maksimovic et al., 2022). On the noisy batches, deMULTIplex2 was able to achieve F scores of about 0.8 with a significant lead over other methods. Similarly, on the lung cell line MULTI-seq dataset and PBMC SCMK dataset, deMULTIplex2 was able to correctly retrieve many more singlets compared to other methods.

Of these datasets, the 8-donor SCMK PBMC dataset was of particular interest to us because the authors labeled the cells with both ADTs from single-cell multiplexing kit (SCMK) reagents (BD Biosciences), and the MULTI-seq LMOs (McGinnis et al., 2021). The authors observed classification of the MULTI-seq tags was much better than classification of the SCMK tags, with the latter showing cell-type biases. We asked if deMULTIplex2 can classify more genuine singlets despite this technological and biological bias. When applying deMULTIplex2 to this dataset, we were able to reproduce the authors’ observation that classification on the MULTI-seq tag count resulted in much better results compared to the SCMK results (Fig. 3B). Compared to previous deMULTIplex-based classification and results from other classification methods, deMULTIplex2 was able to achieve higher recall on the noisy SCMK dataset with high precision (Fig. 3C,D). BFF_cluster_ was able to achieve comparable precision and recall on six donors, but the method performs very poorly on the other two donors with noisy tag data (Fig. 3D). Among these methods, demuxmix also generates probabilistic assignment like deMULTIplex2, allowing us to compare the Receiver Operating Characteristic (ROC) between the two methods. As shown in Fig. 3E, deMULTIplex2 has a much higher area under the ROC curve (AUROC) compared to the two modes of demuxmix, suggesting this mechanism-guided model better captures the difference between positive and negative cell distributions. Looking at the transcriptomic space, we found cell type bias is still present with deMULTIplex2 classification, but more cells were recovered compared to the deMULTIplex result (Fig. 3F,G). However, when comparing to other results from existing tools, deMULTIplex2 generates much lower cell-type bias, and significantly higher classification accuracy (Fig. 3G).

### deMULTIplex2 can salvage cells from complex experiments using precious samples

Many single cell experiments are carried out on precious samples with limited source material, such as tumor cells from patients (Rozenblatt-Rosen et al., 2020) or rare cell populations during development (Zhu et al., 2020). Using single cell multiplexing technology on these samples can reduce batch effects, but depending on the sample quality and cell number, the final cell count recovered from each sample may exhibit large variability. Therefore, demultiplexing methods should be able to robustly handle experimental design where the total cell number per sample is variable and maximally salvage cells from low-cell-count samples.

To understand tumor metastasis in breast cancer, Winkler et al. performed MULTI-seq on a large panel of patient-derived xenograft models (PDX) of human breast cancer (Winkler et al., 2022). The collection and sequencing of the tumors were done across three batches, with the tumors being too heterogeneous to be easily separated in gene expression space (Fig. 4A). The three batches comprise multiple samples of varying total cell numbers, with some samples having very few tagged cells. When applying existing demultiplexing methods on this dataset, several methods, including demuxmix, GMM-Demux, and HTODemux, report errors on one or all of the batches, likely due to the samples with low cell number. Although the rest of the methods were able to classify all three batches without reporting errors, their performance was inferior to deMULTIplex2 (Fig. 4B). In the end, deMULTIplex2 was able to correctly retrieve the highest number of real singlets (63.2% of all cells) compared to deMULTIplex (50.8% of all cells), demonstrating an approximate 25% performance increase. Notably, deMULTIplex2 was able to recover two PDX samples that were almost completely missed by deMULTIplex, GMM-Demux and BFF (Fig. 4C). Examining the tag count distribution of one of such sample (HCI011 tumor tagged with tag “Bar2”), we did not observe a clear bimodal distribution, which likely contributes to failure for methods that rely on such an assumption (Fig. 4D). However, in the axis of cosine similarity, the true positive cells and negative cells are well separated with a much more apparent bimodal distribution (Fig. 4D). With the two GLM-NBs fitted in the two separate spaces, deMULTIplex2 was able to correctly recover 70% of HCI011 tumor cells from the sample (the batch contains another HCI011 sample with a different tag, so the actual recall may be even higher) (Fig. 4E). Finally, when checking the RQRs, we found the distribution of predicted negative cells is close to normal but has a heavy right tail, similar to what we observed with noisy, simulated data. The majority of the predicted positive cells deviate significantly from the negative cell distribution in the residual plot, and there are few cells with ambiguous posterior probability near 0.5 (Fig. 4F).

**Figure 4.**
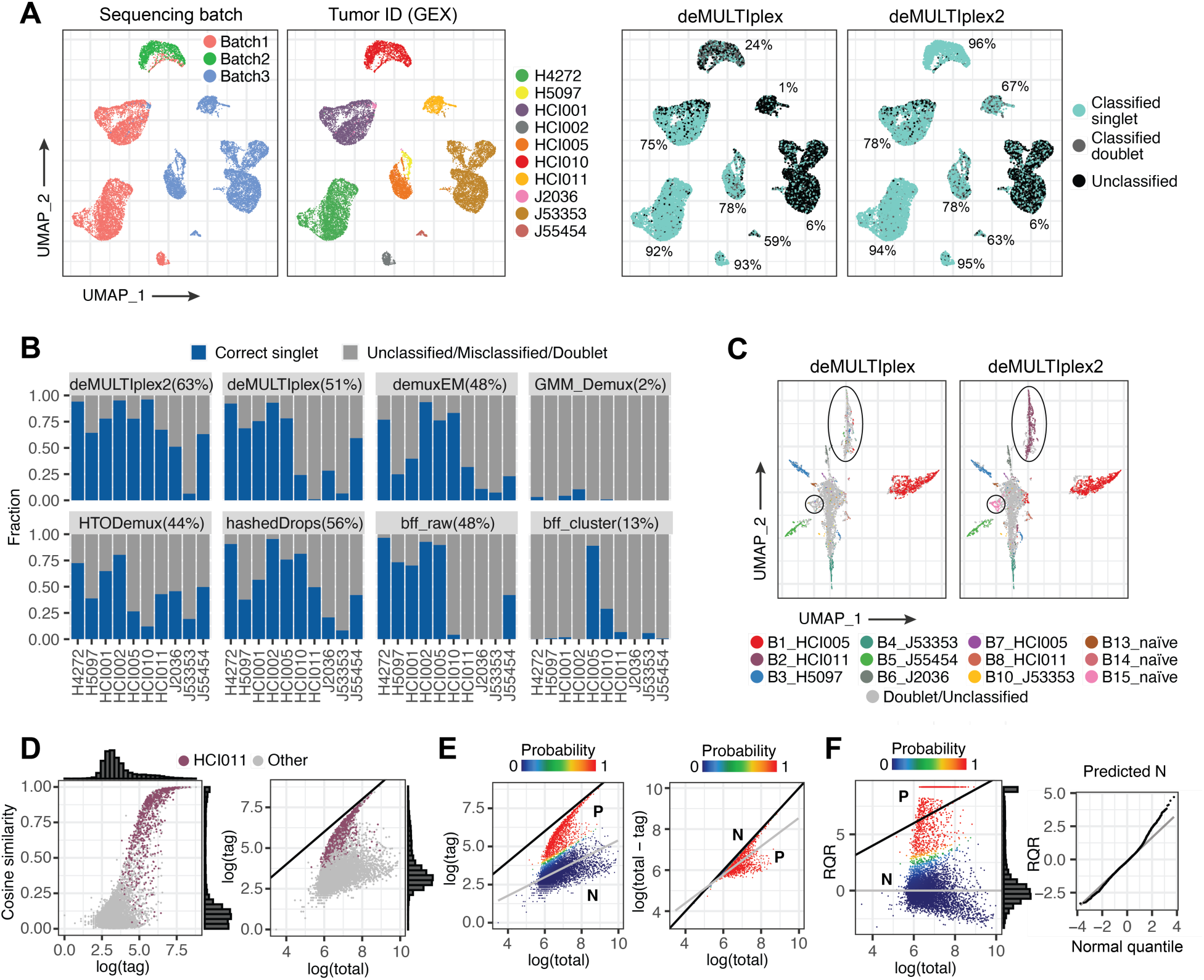
Performance of deMULTIplex2 on multiplexed PDX of human breast cancer. (A) UMAP computed with gene expression (GEX) colored by sequencing batch, expression-based tumor ID, and deMULTIplex or deMULTIplex2 predicted tumor IDs. For major tumor clusters, we highlight the percentage of correctly classified singlets by deMULTIplex and deMULTIplex2, also shown in B. (B) Fraction of cells correctly predicted by deMULTIplex2 and other methods for each tumor model. All methods were run with default parameters. Demuxmix were excluded from the comparison because it returned errors on all three batches. (C) UMAP of cells from batch 3 computed using raw tag UMI counts. Circles highlight two samples that were missed by deMULTIplex with default settings, but were recovered with deMULTIplex2. (D) Cosine similarity vs. tag count plot and the tag count vs. total tag count plot for the sample tagged with “Bar2” and from tumor “HCI011” recovered by deMULTIplex2. The y=x line is shown in black. (E) GLM-NB fit (grey line) and posterior probability of cells being positively tagged by tag “Bar2” calculated by deMULTIplex2 in the two modeling spaces. The y=x line is shown in black, and majority of negative cells fall on or near that line in the second space. N: negative cells, P: positive cells. (F) RQR plotted against log total tag count, colored by posterior probability. For some positive cells, its RQR is infinity. These values were capped to the maximum value of non-infinity RQRs plus 1 for visualization purposes. N: negative cells, P: positive cells. The Q-Q plot compares the distribution of RQRs of predicted negative cells to that of a normal distribution.

Thus, deMULTIplex2 can properly handle complex multiplexed scRNA-seq experiments with precious samples, recover noisy sample tags from these experiments, and does not require the users to pre-filter the tags based on cell number and sample quality. By doing so, it significantly improves the quality of downstream analyses that depend on cell number, such as differential gene expression analysis.

### Improvements on speed and memory

The deMULTIplex package was designed as a complete demultiplexing pipeline which starts from the preprocessing of raw tag FASTQ files (McGinnis, Patterson, et al., 2019). In deMULTIplex2, we have overhauled the code to improve preprocessing steps.

Specifically, deMULTIplex2 utilizes the sparse matrix data structure to efficiently tabulate the tag count of each cell, greatly accelerating computation and reducing the required memory.

As shown previously, the classification algorithm of deMULTIplex2 is highly robust even with down-sampling of cells. Therefore, when running on large single cell datasets, deMULTIplex2 performs down-sampling by default when fitting the GLM-NBs. A 100,000 single cell dataset can be classified by deMULTIplex2 on a MacBook in only a couple of minutes. The software also outputs publication-quality summary and diagnostic plots for users to examine the results in detail.

## Discussion

Although many existing demultiplexing methods work well on clean and small multiplexed datasets, their performance deteriorates rapidly when processing data with large numbers of samples and with noise arising from cross-contamination of tags. Such datasets have become quite common with the increased popularity of single cell multiplexing technologies. This motivated us to develop deMULTIplex2, which is built on a statistical model of tag count distributions derived from the physical mechanism of the contamination process. We found that by modeling contamination from two sources, the cell-bound contamination and the ambient contamination, the observed tag distribution from many real-world experiments can be recapitulated. Using generalized linear models and EM, we were able to probabilistically infer the sample identity of each cell and classify each cell with high confidence. Using real and simulated datasets, we demonstrated that deMULTIplex2 significantly outperforms other methods in recovering genuinely-tagged singlets without compromising precision. This improvement in performance is particularly valuable for real-world applications, as many multiplexed scRNA-seq experiments are carried out on precious samples with limited number of starting cells. More broadly, real world datasets often suffer from higher background and barcode variability that hinders sample classification using previously reported algorithms. Methods that were able to achieve high precision often fail to recover the majority of the true singlets, because they are only classifying cells with the highest signal-to-noise ratio. Similarly, methods that have higher cell recovery often have lower precision, as they are misclassifying cells with noisy tag signals. deMULTIplex2 enables recovery of significantly more cells from these samples without sacrificing classification accuracy, as it is able to correctly draw the decision boundary to separate true positive cells and negative cells. Encouragingly, this behavior is seen consistently across multiple datasets of various cell types generated with distinct multiplexing technologies, suggesting the two-source contamination model broadly captures the tag distribution from these experiments.

deMULTIplex2 is built on several modern statistical techniques with the EM algorithm and generalized linear model at its core. Although the EM algorithm is well known to be susceptible to local optima, we found it was surprisingly robust in deMULTIplex2, even when down-sampling cells to speed up the processing on large-scale datasets. The robustness in performance is achieved through deMULTIplex2’s unique modeling of tag count distributions in two separate spaces. In the first space, we model the observed contaminating tag count of negative cells as a function of total tag count, which is approximately linear in log scale. For positive cells, the tag count quickly converges to the y=x line with increased total tag count and signal-to-noise ratio, and is non-linear when ambient contamination is high. Therefore, the positive cells and the negative cells follow distinct distributions. Similarly, in the second space where we model the total contaminating tags of positive cells as a function of total tag count, the negative cells converge to the y=x line with increased total tag count, which significantly deviates from the positive cells. By fitting two GLM-NB models in these separate spaces, deMULTIplex2 is able to discriminate these distinct distributions of positive and negative cells, leading to increased accuracy and robustness. When diagnosing the model fit with Randomized Quantile Residuals, we found the RQRs approximately follow the normal distribution, suggesting the fitted model was able to explain the variance in the observed tag count. Finally, we provide a cosine-similarity-based initialization for the EM algorithm, which by itself better captures the bimodal distribution of tags compared to raw tag UMI counts and further improves the robustness of the algorithm.

In the deMULTIplex2 algorithm, we assume a simple generalized linear model. We base this assumption on the two-source contamination model, in which the log of expected contaminating tag counts is linear with ln(*N_total_* + *C*) as a predictor, assuming total tag counts of most cells are within the range of one or two orders of magnitude. Although this is indeed true on many real datasets we tested, the model may work suboptimally on highly heterogenous datasets with a wide range of cell size. Therefore, more sophisticated modeling, such as a better estimation of *C* – perhaps by utilizing tag counts from empty droplets – may generate even better classification performance. Another limitation of deMULTIplex2 is doublet calling, as its current doublet detection capability is not significantly better than existing methods. This is likely because deMULTIplex2 does not explicitly model the distribution of doublets. Therefore, we still recommend users to run other doublet detection tools, such as DoubletFinder (McGinnis, Murrow, et al., 2019) and Scrublet (Wolock et al., 2019), to remove heterotypic doublets.

In summary, deMULTIplex2 models the physical process of sample tag cross-contamination to correctly assign sample-of-origin in real world sample multiplexing experiments. By applying generalized linear models and expectation-maximization, deMULTIplex2 can achieve significant performance improvement on both simulated and real-world datasets compared to existing algorithms. deMULTIplex2 helps users salvage genuine singlet cells from the non-idealized conditions encountered in real world experiments without sacrificing classification accuracy, thus greatly improving the quality of downstream analysis.

## Methods

### Derivation of the contamination model

We model the contamination of tag counts as two components: the contamination of cell-bound tags, which is correlated with cell surface area, and the contamination from ambient floating tags captured by the droplet, which correlated with droplet size and assumed to be constant across all cells. For a particular tag B, cells can be divided into two partitions – those “positively” labeled with tag B before pooling, and those that should be “negative” for B but got contaminated by B after pooling. For each negative cell, if we denote total observed tag count as *N_total_*, the ambient tag counts for each tag k as *M_k_*, and assume cell-bound tag count of B takes a constant fraction *p_B_* of total cell bound tag 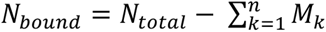, then the expected contamination level of B on a negative cell can be written as

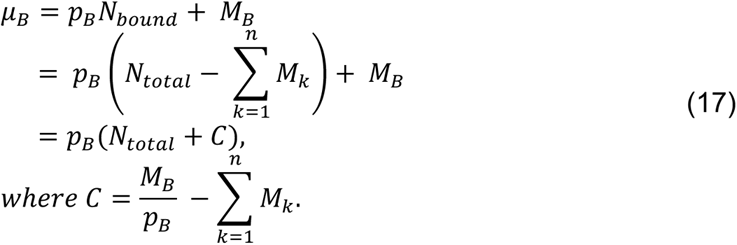

Taking the log transform of *μ_B_* we can obtain the following equation:

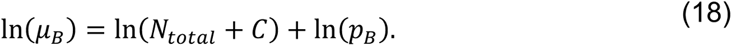

Following the argument made in the result section, we approximate the above equation with

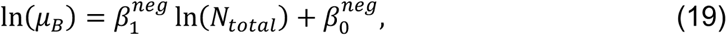

Assuming the observed count follows a negative binomial distribution:

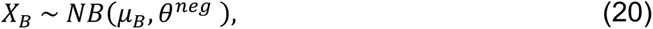

then a negative binomial generalized linear model can be applied to the tag count of negative cells to estimate the parameters 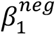, 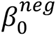, *θ^neg^*.

For positive cells originally tagged with B, tag count B will be equal to the total tag count under ideal conditions, but often deviates from the y=x line due to contamination (Fig. 1, S1). As discussed previously, we choose to model the “contamination part” of the positive cells because the contamination of all cells happened after pooling and can be modeled uniformly. Equation 17 shows for a single contaminating tag B, its expected count follows:

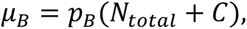

Then for a pool of contaminating tags,

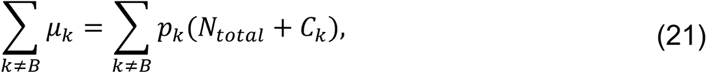

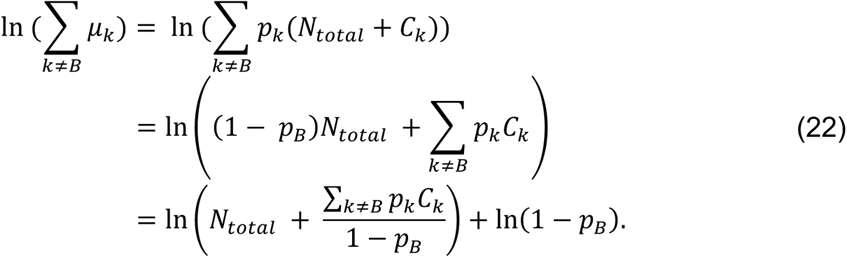

This result is in a form similar to equation 18, therefore we can use the approximation below when fitting a GLM-NB model:

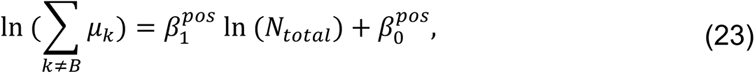

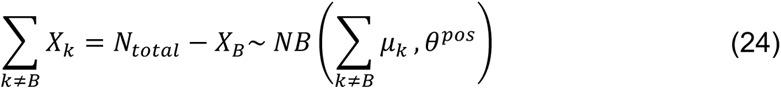

### Implementation of expectation-maximization (EM)

We implemented EM using the R programming language. We first initialize the algorithm from the M step with a non-random guess based on the cosine similarity with the canonical vector of each tag. The method then iterates between the E step and the M step to maximize the joint log likelihood until convergence or when the maximal number of iterations has been reached.

In the M step, we fit the GLM-NB on the negative cells in the *X* vs *N_total_* space, and the positive cells in the *N_total_* − *X* vs *N_total_* space with the log link function, i.e.,

For negative cells:

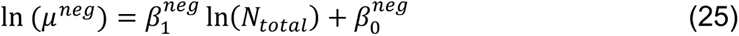

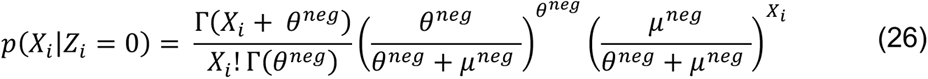

For positive cells:

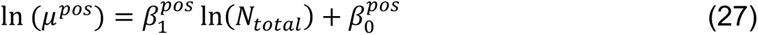

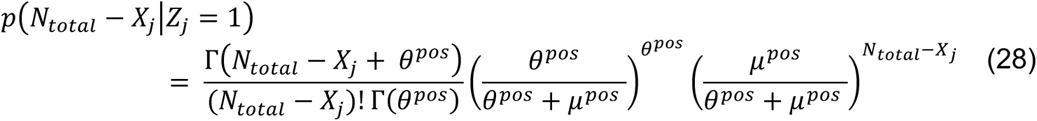

The model parameters, 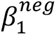, 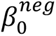, *θ^neg^* and 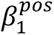, 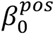, *θ^pos^* are estimated with the *glm.nb* function from the *MASS* package.

In the E step, the posterior probabilities of the cells are calculated based on the GLM-NB predicted probability and prior probability, i.e.,

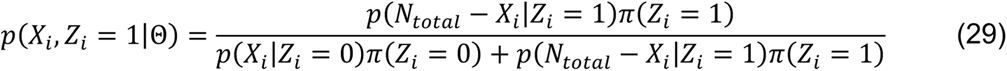

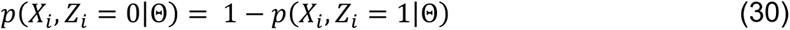

The prior probability, *π*(*Z_i_* = *0*) and *π*(*Z_i_* = 1) are defined as the fraction of the cells being negative or positive given the posterior probability from previous iteration with a 0.5 cutoff.

In the *X* vs *N_total_* space, *p*(*X_i_*|*Z_i_* = *0*) estimated from the GLM-NB is highest around the fitted mean and becomes lower when the observed tag count deviates from the mean. However, cells with tag count lower than the fitted mean are theoretically more likely to be negative cells. Therefore, when calculating the posterior probability, we set *p*(*X_i_*|*Z_i_* = *0*) of cells below the mean to be the same as those estimated at or near the mean when computing the posterior probability, i.e.,

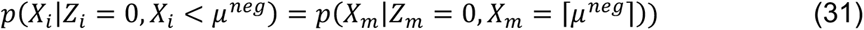

This adjustment allows more robust classification performance. In addition, we found that when fitting the model with all cells at the M step, the algorithm typically converges in a few iterations. However, with very large dataset, fitting the GLM-NB model with all cells can be computationally expensive. In practice, we down-sample the positive and negative cells to some user-specified number to expediate the fitting process. We found that when using a reasonable number of down-sampled cells to fit the model, the algorithm quickly and robustly converges in a limited number of iterations (Fig. S2).

### Generating simulated datasets

To simulate realistic tag count data, we took a three-step approach to reproduce the labeling, pooling, and contamination process *in silico*. In the first step, we sample from a user-defined range to generate the initial tag count, which is a clean matrix with only positive cells having non-zero entries for the corresponding tag (denoted as *X_True_*). In the second step, we assume that the initial count is correlated with cell surface area, and use this vector as the total tag count *N_total_* in equation 2 to generate the expected cell-bound contamination count *μ_B_* with user specified *p_B_* for each tag. We then sample counts from a negative binomial distribution with the cell-specific expected contamination level *μ_B_* and user-defined overdispersion parameter *θ_B_* to generate the cell-bound contamination matrix (denoted as *C_cell-bound_*). Finally, to simulate the ambient contamination, we sample from a user-defined range to obtain the expected ambient level for each tag. We then perform negative binomial sampling to generate realistic noise for each tag, and obtain an ambient contamination matrix *C_ambient_*. The final singlet count matrix is generated by summing up the three matrices *X_True_*, *C_cell-bound_* and *C_ambient_*. To generate doublets, we randomly sample pairs of singlets up to a user-defined percentage, and sum up the corresponding entries in *X_True_* and *C_cell-bound_*. Because doublets are generally encapsulated within a single droplet, we only sample *C_ambient_* once and add the value to the simulated doublets.

For the simulated data displayed in Fig. 2, we sample from 10 to 1000 to define the expected value of true tag count, and from 0 to 100 to define the expected value of ambient contamination. The overdispersion parameter of the NB distribution *θ* was set to 10, which is in the range of estimated *θ* from real datasets (Fig. S1A). We set ln (*p_B_*) = −4 for all contaminating tags, which means the cell-bound contaminating tag takes up about 1.8% of total cell-bound tags, representing a mild contamination level. Finally, for both 5-tag simulation and 30-tag simulation, we set the doublet rate to 10%, which is also common in droplet-based single cell experiments.

### Benchmarking on real datasets

To prepare public datasets for benchmarking, we preprocessed the tag count matrix from each of the studies into a uniform format and include their ground truth labeling when available. For multi-donor datasets, we ran SNP-based sample classification using vireo (Huang et al., 2019) or souporcell (Heaton et al., 2020) when such genotype-based classification were not provided.

All the methods we benchmarked were run using their default parameter setting without any parameter tuning. demuxEM and demuxmix are two methods that require information from the transcriptome. Thus, we were not able to benchmark these methods using the simulated datasets or with datasets which did not provide such information. deMULTIplex2 was also run with default parameters across all benchmarking cases, with initial cosine cut off set to 0.5, max number of cells for GLM-NB fitting set to 5000, and max number of EM iterations set to 30.

## Acknowledgements

We thank the members of the Gartner Lab for sharing datasets, providing critical discussions and support. We also thank members of the Matt Thomson Lab at Caltech for sharing datasets for testing deMULTIplex2.

## Funding

This research was supported by grants from the NIH (U01CA199315, R01GM135462, R01DK126376, U01DK103147, and R33CA247744) and the UCSF Center for Cellular Construction (DBI-1548297), an NSF Science and Technology Center. Z.J.G. is a Chan Zuckerberg BioHub San Francisco Investigator.

## Availability of data and materials

Demultiplex2 is available as an R package at https://github.com/Gartner-Lab/deMULTIplex2 under the Creative Commons Attribution-NonCommercial-NoDerivatives 4.0 International License (http://creativecommons.org/licenses/by-nc-nd/4.0/).

## Contributions

Q.Z., D.N.C. and Z.J.G. conceived the project. Q.Z. designed and implemented the algorithm. Q.Z. and D.N.C. implemented preprocessing and plotting functions of the R package and documented the R package. Q.Z. carried out benchmarking analysis with help from D.N.C. Z.J.G. supervised the project. Q.Z. and Z.J.G. wrote the manuscript. All authors reviewed and edited the manuscript.

## Competing interests

ZJG is an author on patents associated with sample multiplexing and ZJG is an equity holder and advisor to Scribe Biosciences and Serotiny.

## Supplemental Figures

**Figure S1.**
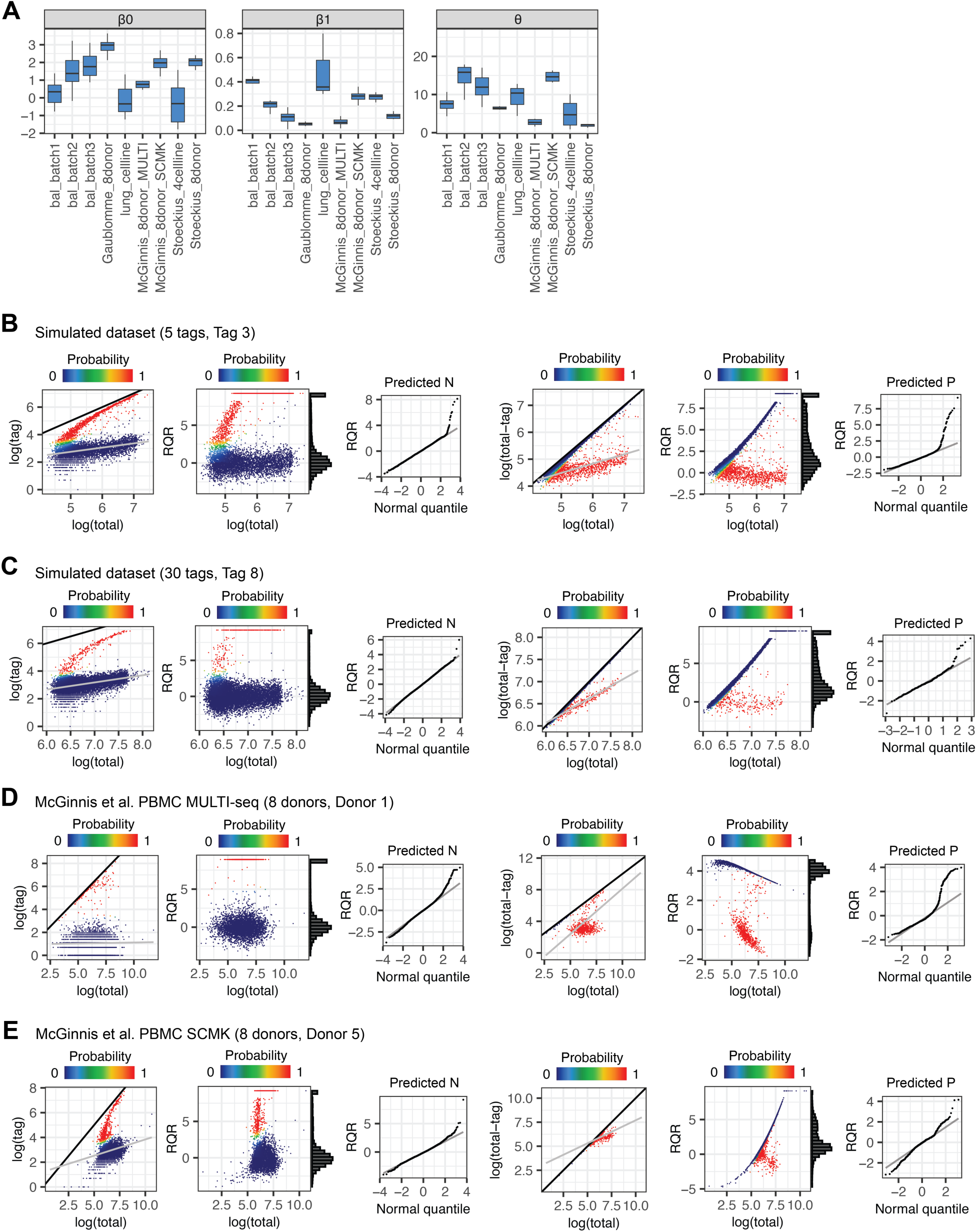
deMULTIplex2 estimated parameters and examples of tag count distribution. (A) deMULTIplex2 estimated parameters of the GLM-NB model of negative cells for tags across 9 real datasets. (B-E) For each dataset, a random tag is picked and its count distributions are plotted in the two modeling spaces, colored by the deMULTIplex2-inferred posterior probability of the cells being positively tagged. The y=x line is shown in black, and majority of negative cells fall on or near that line in the second space. RQRs are computed for both GLM-NB fits and plotted against the normal quantiles. For some cells, its RQR is infinity. These values were capped to the maximum value of non-infinity RQRs plus 1 for visualization purpose.

**Figure S2.**
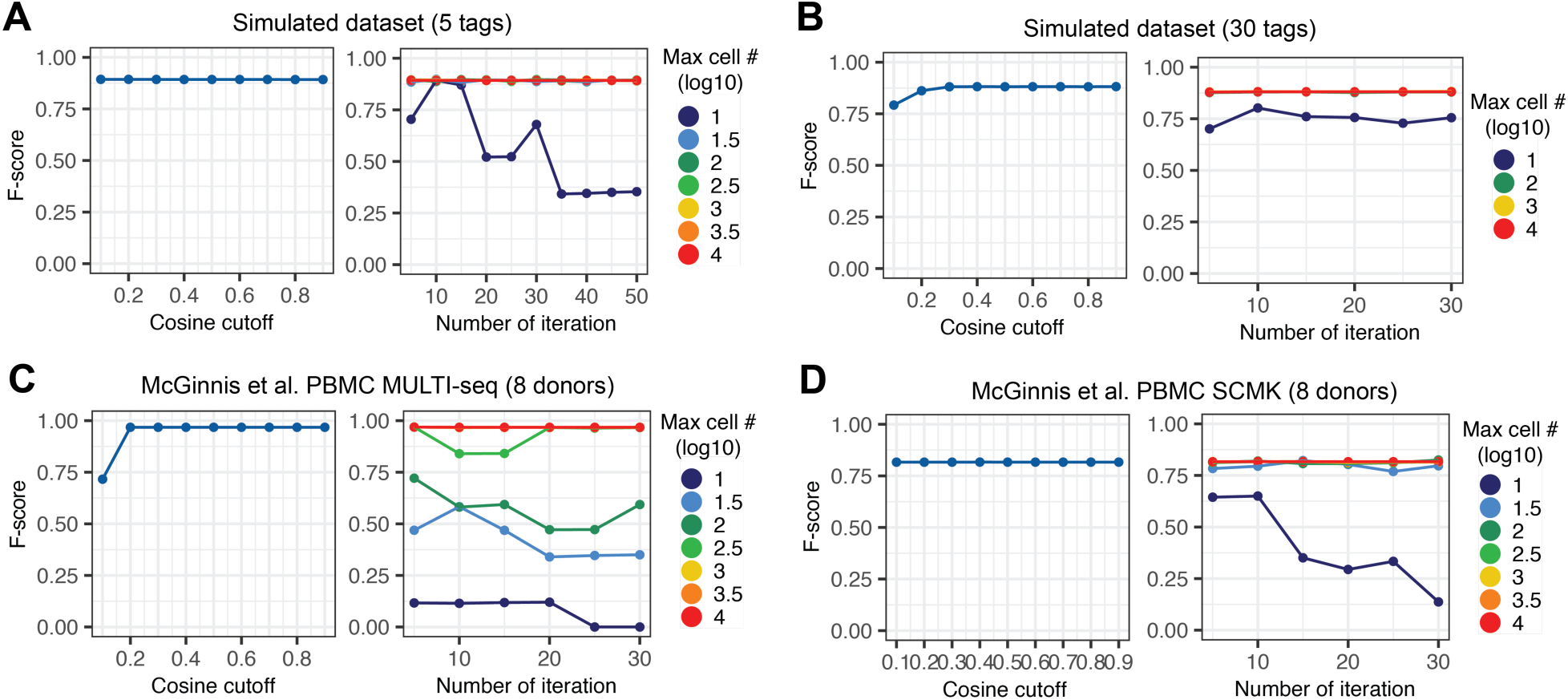
Robustness of deMULTIplex2 against parameter values and down-sampling. (A-D) Performance of deMULTIplex2 (measured by average F score) on the two simulated datasets presented in Fig. 2, and the MULTI-seq and SCMK datasets from McGinnis et al. presented in Fig. 3 (McGinnis et al., 2021). The left panel shows the performance of deMULTIplex2 when the EM algorithm is initialized with positive and negative cells determined by various cutoffs on cosine similarity. The right panel shows the performance of deMULTIplex2 when the GLM-NBs were fit with down-sampled positive and negative cells and the indicated number of iterations of the EM algorithm.

**Figure S3.**
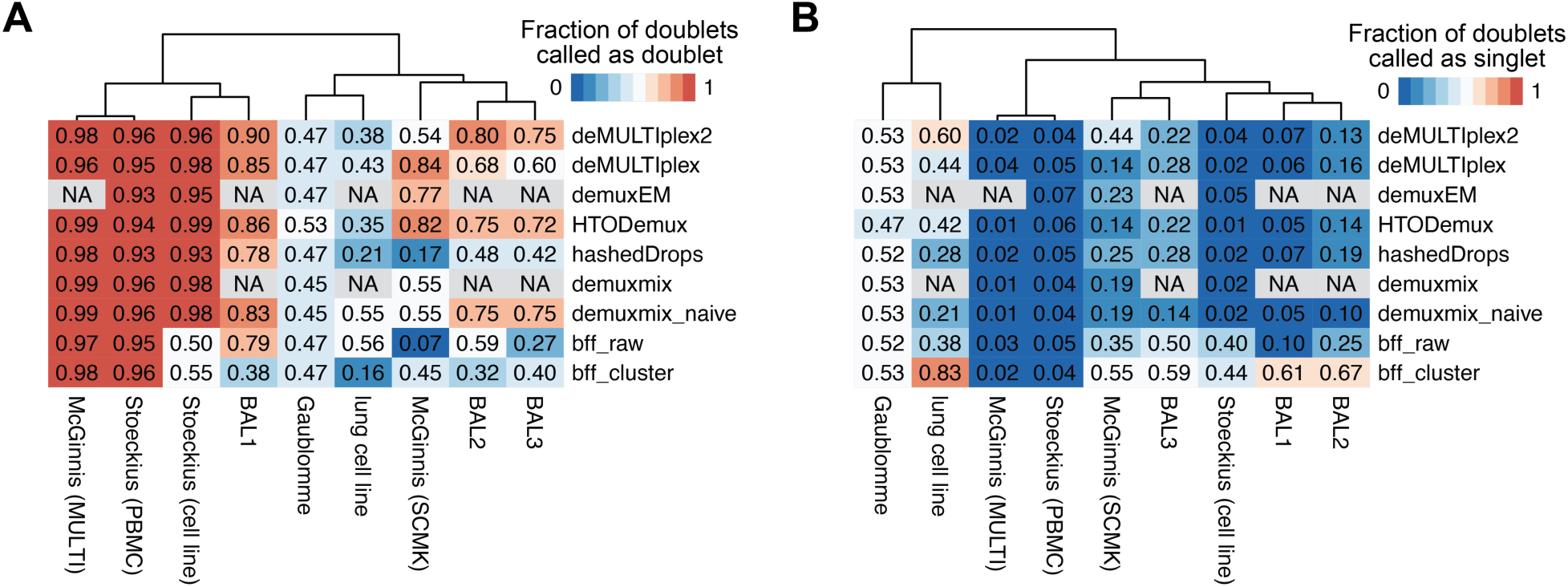
Doublet calling of deMULTIplex2 and other methods on real datasets. (A) Heatmap showing the fraction of true doublets called correctly by deMULTIplex2 and existing methods. (B) Heatmap showing the fraction of doublets misclassified as singlets by these methods. NA indicates the methods cannot be run on the corresponding datasets due to the unavailability of an mRNA count matrix or error (i.e., demuxEM returns an error on the SCMK dataset). GMM-Demux was excluded from this analysis because the method only generates singlet classifications.

## Supplemental Tables

**Supplemental Table 1. Performance of deMULTIplex2 and other methods on real datasets.** For each method, precision, recall, and F score are reported for each dataset. Empty values indicate the method cannot be run on the corresponding dataset due to unavailability of mRNA count matrix or an error.

## Notes

### Summary of Updates

Title corrected from Demultiplex2 to deMULTIplex2. Author name corrected from Danny N. Conrad to Daniel N. Conrad.

